# Ultra-deep, long-read nanopore sequencing of mock microbial community standards

**DOI:** 10.1101/487033

**Authors:** Samuel M. Nicholls, Joshua C. Quick, Shuiquan Tang, Nicholas J. Loman

## Abstract

**Background:** Long sequencing reads are information-rich: aiding *de novo* assembly and reference mapping, and consequently have great potential for the study of microbial communities. However, the best approaches for analysis of long-read metagenomic data are unknown. Additionally, rigorous evaluation of bioinformatics tools is hindered by a lack of long-read data from validated samples with known composition.

**Methods:** We sequenced two commercially-available mock communities containing ten microbial species (ZymoBIOMICS Microbial Community Standards) with Oxford Nanopore GridION and PromethION. Both communities and the ten individual species isolates were also sequenced with Illumina technology.

**Data:** We generated 14 and 16 Gbp from GridION flowcells and 146 and 148 Gbp from PromethION flowcells for the even and log communities respectively. Read length N50 was 5.3 Kbp and 5.2 Kbp for the even and log community, respectively. Basecalls and corresponding signal data are made available (4.2 TB in total).

**Results:** Alignment to Illumina-sequenced isolates demonstrated the expected microbial species at anticipated abundances, with the limit of detection for the lowest abundance species below 50 cells (GridION). *De novo* assembly of metagenomes recovered long contiguous sequences without the need for pre-processing techniques such as binning.

**Conclusions:** We present ultra-deep, long-read nanopore datasets from a well-defined mock community. These datasets will be useful for those developing bioinformatics methods for long-read metagenomics and for the validation and comparison of current laboratory and software pipelines.

## Data Description

Whole-genome sequencing of microbial communities (metagenomics) has revolutionised our view of microbial evolution and diversity, with numerous potential applications for microbial ecology, clinical microbiology and industrial biotechnology [1, 2]. Typically, metagenomic studies use high-throughput sequencing platforms (*e.g.* Illumina) [3] which generate very high yield, but of limited read length (100–300 bp).

In contrast, single-molecule sequencing platforms such as the Oxford Nanopore MinION, GridION and PromethION are able to sequence very long fragments of DNA (>10 Kbp, with over 2 Mbp reported) [4, 5] and with recent improvements to the platform making metagenomic studies using nanopore more viable, such studies are increasing in frequency [6, 7, 8, 9]. Long reads help with alignment-based assignment of taxonomy and function due to their increased information content [10, 11]. Additionally, long reads permit bridging of repetitive sequences (within and between genomes), aiding genome completeness in *de novo* assembly [12]. However, these advantages are constrained by high error rate (≈10%), requiring the use of specific long-read alignment and assembly methods, which are either not specifically designed for metagenomics, or have not been extensively tested on real data [13].

Mock community standards are useful for the development of genomics methods [14], and for the validation of existing laboratory, software and bioinformatics approaches. For example, validating the accuracy of a taxonomic identification pipeline is important, because the consequences of erroneous taxonomic identification from a metagenomic analysis may be severe, *e.g.* in public health microbiology (such as in the well-publicised case of anthrax and plague in the New York Subway [15]) or missed diagnoses in the clinic. Mock community standards can also be used as positive controls during laboratory work, for example to validate that DNA extraction methods will yield expected representation of a sampled community [14].

Here, we present four nanopore sequencing datasets of two microbial community standards, providing a state-of-the-art benchmark to accelerate the development of methods for analysing long-read metagenomics data.

### Background Information

The ZymoBIOMICS Microbial Community Standards (CS and CSII) are each composed of ten microbial species: eight bacteria and two yeasts (Table 1). The organisms in CS (hereafter referred to as ‘Even’) are distributed equally (12%), with the exception of the two yeasts which are each present at 2%. Cell counts from organisms in CSII (‘Log’) community are distributed on a log scale, ranging from 89.1% (*Listeria monocytogenes*), down to 0.000089% (*Staphylococcus aureus*).

## Methods

### DNA extraction

DNA was extracted from 75 µl ZymoBIOMICS Microbial Community Standard (Product D6300, Lot ZRC190633) and 375 µl ZymoBIOMICS Microbial Community Standard II (Product D6310, Lot ZRC190842) using the ZymoBIOMICS DNA Miniprep extraction kit according to manufacturer’s instructions, with the following modifications to increase fragment length and maintain the expected representation of the Gram-negative species which are already lysed in the DNA/RNA Shield storage solution. The standard was spun at 8,000×*g* for 5 minutes before removing the supernatant and retaining. The cell pellet was resuspended in 750 µl lysis buffer and added to the ZR BashingBead lysis tube. Bead-beating was performed on a FastPrep-24 (MP Biomedicals) instrument for 2 cycles of 40 seconds at power level 6, with 5 minutes sitting on ice between cycles. The bead tubes were spun at 10,000×*g* for 1 minute and 450 µl of supernatant was transferred to a Zymo Spin III-F filter before being spun again at 8000×*g* for 1 minute. 45 µl (Even) and 225 µl (Log) of the supernatant retained earlier was combined with 450 µl filtrate before adding 1485 µl (Even) or 2025 µl (Log) Binding Buffer and mixing before loading onto the column.

**Table 1.**
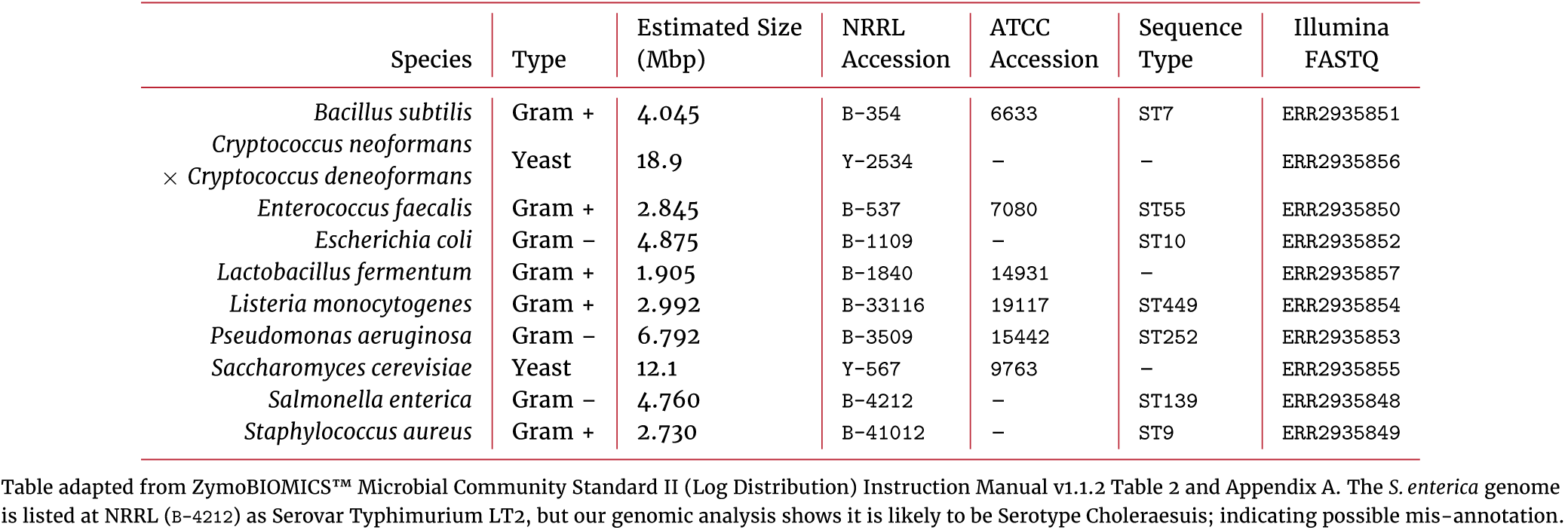
Description of the ten organisms comprising the ZymoBIOMICS Mock Community Standards.

### Nanopore sequencing library preparation

Quantification steps were performed using the dsDNA HS assay for Qubit. DNA was size-selected by cleaning up with 0.45× volume of Ampure XP (Beckman Coulter) and eluted in 100 µl EB (Qiagen). Libraries were prepared from 1400 ng input DNA using the LSK-109 kit (Oxford Nanopore Technologies) as per manufacturer’s protocol, except incubation times for end-repair, dA-tailing and ligation were increased to 30 minutes to improve ligation efficiency. The even and log libraries were split and used on both the GridION and PromethION flowcells.

### Sequencing

Sequencing libraries were quantified and two aliquots of 50 ng and 400 ng were prepared for GridION and PromethION sequencing respectively. The GridION sequencing was performed using a `FLO-MIN106` (rev.C) flowcell, `MinKNOW` 1.15.1 and standard 48-hour run script with active channel selection enabled. The PromethION sequencing was performed using a `FLO-PRO002` flowcell, `MinKNOW` 1.14.2 and standard 64-hour run script with active channel selection enabled.

Refuelling was performed approximately every 24h (GridION, PromethION) by loading 75 µl (GridION) or 150 µl (PromethION) refuelling mix (SQB diluted 1:2 with nuclease-free water). Additionally, after the standard scripts had completed the PromethION was restarted several times to utilise remaining active pores and maximise total yield.

### Nanopore basecalling

Reads were basecalled on-instrument using the GPU base-caller `Guppy` (Oxford Nanopore Technologies) with the supplied `dna_r9.4.1_450bps_prom.cfg` configuration (PromethION) and `dna_r9.4_450bps.cfg` (GridION). `Guppy` version 1.8.3 and 1.8.5 were installed on the GridION and PromethION respectively.

### Illumina sequencing

DNA was extracted from pure cultures of each species using the ZymoBIOMICS DNA Miniprep Kit. Library preparation was performed using the Kapa HyperPlus Kit with 100 ng DNA as input and TruSeq Y-adapters. The purified library derived from each sample was quantified by TapeStation (Agilent 4200) and pooled together in an equimolar fashion. The pooled library was sequenced on an Illumina HiSeq 1500 instrument using 2×101 bp (paired-end) sequencing, over four lanes. Raw reads were demultiplexed using `bcl2fastq` v2.17. Shotgun sequencing of the even and log communities was performed with the same protocol, with the exception that the log community was sequenced individually on two flowcell lanes, and the even community was instead sequenced on an Illumina MiSeq using 2×151 bp (paired-end) sequencing.

### Bioinformatics

To construct reference genomes, Illumina reads for each of the ten isolates were assembled using `SPAdes` v3.12.0 [16] with paired-end reads as input, using parameters `-m 512–t 12`. Scaffolds from `SPAdes` less than 500 bp length or with less than 10× coverage were removed. The remaining scaffolds were combined into a single mock community draft assembly for downstream analysis. Multilocus sequence typing (MLST) of the scaffolds was conducted with `mlst` (https://github.com/tseemann/mlst), using default parameters.

Nanopore reads were aligned to the mock community draft assembly using `minimap2` [17] v2.14-r883 with parameters `-ax map-ont -t 12` and converted to a sorted BAM file using `samtools` [18]. To reduce erroneous mappings, alignment BAM files were filtered using a script `bamstats.py` according to the following criteria; reference mapping length ≥500 bp, map quality (MAPQ) > 0, there are no supplementary alignments for this read and read is not a secondary alignment. Per-species coverage summary statistics were generated using the `summariseStats.R` Rscript.

Read accuracy was determined by calculating `BLAST` identity from the filtered alignments (as per http://lh3.github.io/2018/11/25/on-the-definition-of-sequence-identity), calculated as (*L*–*NM*)/*L* using the `minimap2` number of mismatches (`NM`) SAM tag and the sum of match, insertion and deletion CIGAR operations (L).

We used `wtdbg2` v2.2 to assemble nanopore data (https://github.com/ruanjue/wtdbg2). `wtdbg2` was compiled from source in November 2018 from Git commit `094f2b3`. For GridION, all nanopore reads were used. For PromethION, we used both the full dataset and a random 25% subsample. Subsampling was performed using `subsample.py`.

Assemblies were conducted under a variety of parameter values for homopolymer-compressed k-mer size (`-p`), minimum graph edge weight support (`-e`) and read length threshold (`-L`). Global parameters for all runs (`-S1 -K10000 -node-max 6000`) were used to turn-off k-mer subsampling (to remove assembly stochasticity) and increase the coverage thresholds applied to k-mers and constructed nodes.

Assembled contigs were assigned to taxa with `kraken2` [19] (`--use-names -t12`) using a database containing all of the archaeal, bacterial, fungal, protozoal and viral sequences from `RefSeq`, and `UniVec_Core` (database download links are in our repository). The `kraken2` output was parsed with `extracken.py`. Assembly accuracy was determined by a modified version of `fastmer.py` (https://github.com/jts/assembly_accuracy) which uses `minimap2` to align contigs against the draft assembly.

## Results

### Nanopore sequencing metrics

We generated a total of 324.36 Gbp of sequence from the four nanopore sequencing runs (Table 2, Figure 3.a). PromethION flowcells generated approximately ten-times more sequencing data than the comparative GridION runs and showed equivalent read length N50 and quality scores (Figure 1.b). We observe a difference in sequencing speed between the PromethION (mean 395 bps and 412 bps for even and log) and the GridION (mean speed 341 and 359 bps for even and log) (Figure 1.c).

**Table 2.** Summary of the four nanopore sequencing experiments.

| Signal Accession | FASTQ Accession | Sequencer | Standard | Time (h) | Reads (M) | N50 (Kbp) | Quality (Median Q) | Yield (Gbp) | Q>7 (Gbp) |
| --- | --- | --- | --- | --- | --- | --- | --- | --- | --- |
| ERR2887847 | ERR2906227 | GridION | Zymo CS Even ZRC190633 | 48 | 3.49 | 5.3 | 9.8 | 14.01 | 12.31 |
| ERR2887850 | ERR2906229 | GridION | Zymo CSII Log ZRC190842 | 48 | 5.73 | 5.2 | 9.3 | 16.03 | 13.67 |
| ERR2887848<br>ERR2887849 | ERR2906228 | PromethION<br>PromethION | Zymo CS Even ZRC190633<br>Zymo CS Even ZRC190633 | 64<br>– | 36.5 | 5.3 | 9.1 | 146.29 | 123.27 |
| ERR2887851<br>ERR2887852 | ERR2906230 | PromethION<br>PromethION | Zymo CSII Log ZRC190842<br>Zymo CSII Log ZRC190842 | 64<br>– | 35.1 | 5.2 | 9.2 | 148.03 | 126.12 |

PromethION runs were restarted following the standard 64 hour protocol. The table reflects total yield across both the standard run and subsequent restarts.

**Figure 1.**
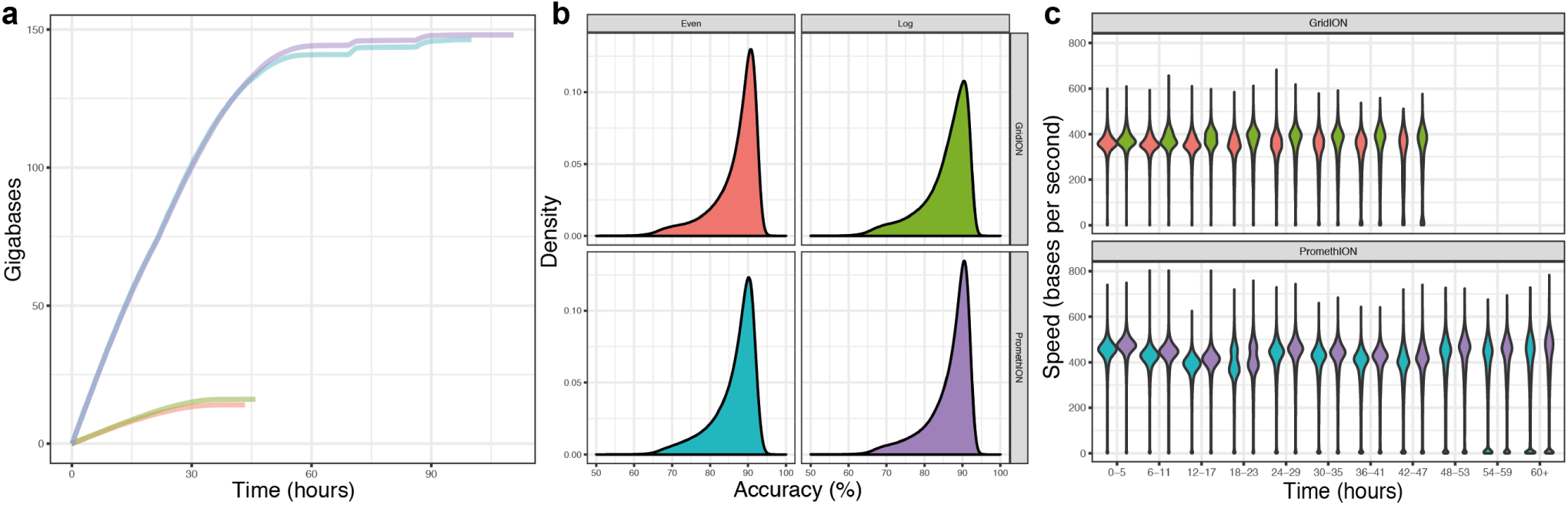
Summary plots for the four generated data sets: (a) collector’s curve showing sequencing yield over time for each of the four sequencing runs, (b) density plot showing sequence accuracy (`BLAST`-like identities), (c) violin plot showing sequencing speed over time split by sequencing group.

**Figure 3.**
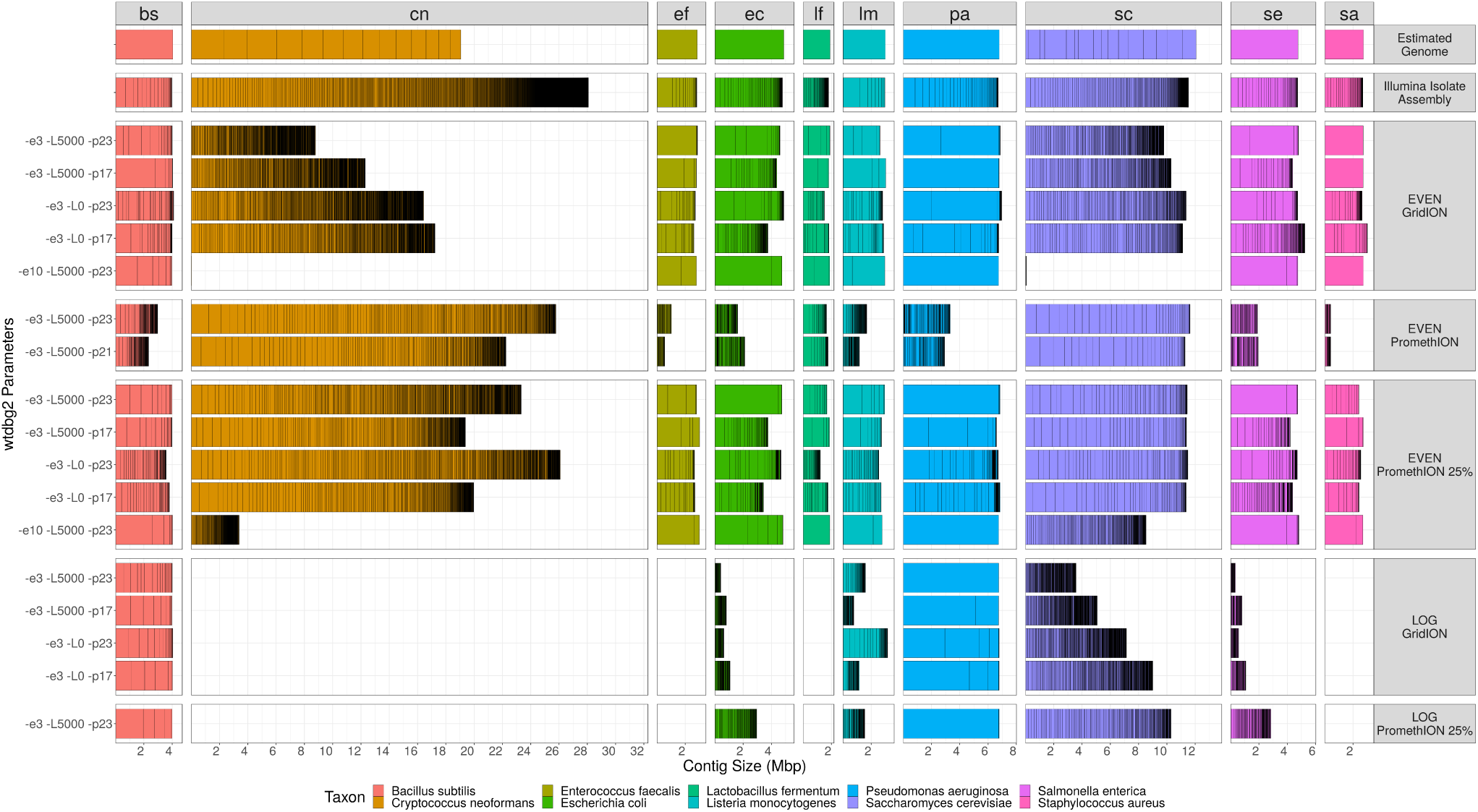
Bar plots demonstrating total length and contiguity of genomic assemblies obtained with `wtdbg2` from each of the long-read nanopore data sets. For each organism in the community (coloured columns), contigs longer than 10 kbp are horizontally stacked along the x-axis. Each row represents a run of `wtdbg2`, with the parameters for edge support, read length threshold and homopolymer-compressed k-mer size labelled on the left. Assemblies are grouped by the data set on which they were run (row facets). Additionally, assemblies may be compared to the estimated true genome size, and per-isolate Illumina `SPADES` assembly. Estimated genomes sizes are the same as those found in Table 1, however to display approximate chromosomes, the two yeasts were replaced by their corresponding canonical NCBI references for visualisation purposes only. The *C. neoformans* strain used by the Zymo standards is a diploid genetic cross, which may explain the larger assemblies, compared to the represented estimated haploid size.

### Illumina sequencing metrics

Illumina datasets for the ten individually sequenced isolates averaged 13.53 million pairs of reads (ranging between 7.1 – 23.2 million), with proportions of reads with a mean `phred` score *≥*30 ranging between 75.51% – 93.09% (Table 5). Illumina sequencing generated 8.8 million pairs of reads (2×151 bp, MiSeq), and 47.8 million pairs of reads (2×101 bp, HiSeq) for the even and log community, respectively (Table 5).

**Table 5.** Summary statistics for Illumina sequencing data.

| Dataset | Pairs (M) | Yield (Gbp) | phred<sub>≥</sub>30 (%) | Accession |
| --- | --- | --- | --- | --- |
| Isolates | 13.53<br>± 5.23 | 2.73<br>± 1.06 | 87.72%<br>± 5.43% | see Table 1 |
| CS (Even) | 8.8 | 2.65 | 95.12 % | ERR2984773 |
| CSII (Log) | 47.8 | 9.66 | 95.71 % | ERR2935805 |

Illumina sequencing was performed on an Illumina HiSeq 1500, with the exception of the even community which was sequenced on an Illumina MiSeq.

### Nanopore mapping statistics

We identify the presence of all 10 microbial species in the community, for both even and log samples, in expected proportions (Figure 2). For the even community, the GridION results provide sufficient depth (i.e. ≫ 30× coverage) to potentially assemble all eight of the bacteria. The coverage of the yeast genomes were lower (10 and 17×), potentially sufficient for assembly scaffolding. On the PromethION all genomes had >100× mean coverage (Tables 3 and 4).

**Table 3.** Read alignment statistics for even samples, showing absolute measurements and proportion of sequencing yield and the estimated genome coverage obtained for each organism in the mock community.

| Species | Expected Proportion | GridION |  |  |  | PromethION |  |  |  |
| --- | --- | --- | --- | --- | --- | --- | --- | --- | --- |
|  |  | Yield (Gbp) | Measured Proportion | Aln. N50 (Kbp) | Coverage (×) | Yield (Gbp) | Measured Proportion | Aln. N50 (Kbp) | Coverage (×) |
| *Bacillus subtilis* | 12 | 2.09 | 19.30 | 4.30 | 516.41 | 21.54 | 18.98 | 4.40 | 5325.31 |
| *Listeria monocytogenes* | 12 | 1.57 | 14.53 | 4.46 | 525.44 | 16.20 | 14.28 | 4.57 | 5414.09 |
| *Enterococcus faecalis* | 12 | 1.32 | 12.21 | 4.45 | 464.33 | 13.66 | 12.04 | 4.56 | 4802.85 |
| *Staphylococcus aureus* | 12 | 1.22 | 11.25 | 4.46 | 445.98 | 12.58 | 11.08 | 4.57 | 4607.07 |
| *Salmonella enterica* | 12 | 1.08 | 10.01 | 8.56 | 227.50 | 11.76 | 10.36 | 8.94 | 2470.85 |
| *Escherichia coli* | 12 | 1.08 | 9.95 | 8.31 | 220.93 | 11.69 | 10.30 | 8.70 | 2397.58 |
| *Pseudomonas aeruginosa* | 12 | 1.05 | 9.73 | 9.00 | 155.08 | 11.58 | 10.20 | 9.37 | 1704.32 |
| *Lactobacillus fermentum* | 12 | 1.01 | 9.30 | 3.6 | 528.46 | 10.35 | 9.12 | 3.72 | 5433.40 |
| *Saccharomyces cerevisiae* | 2 | 0.21 | 1.92 | 4.08 | 17.18 | 2.12 | 1.87 | 4.17 | 174.97 |
| *Cryptococcus neoformans* | 2 | 0.19 | 1.79 | 4.44 | 10.23 | 2.00 | 1.76 | 4.53 | 105.95 |

**Table 4.**
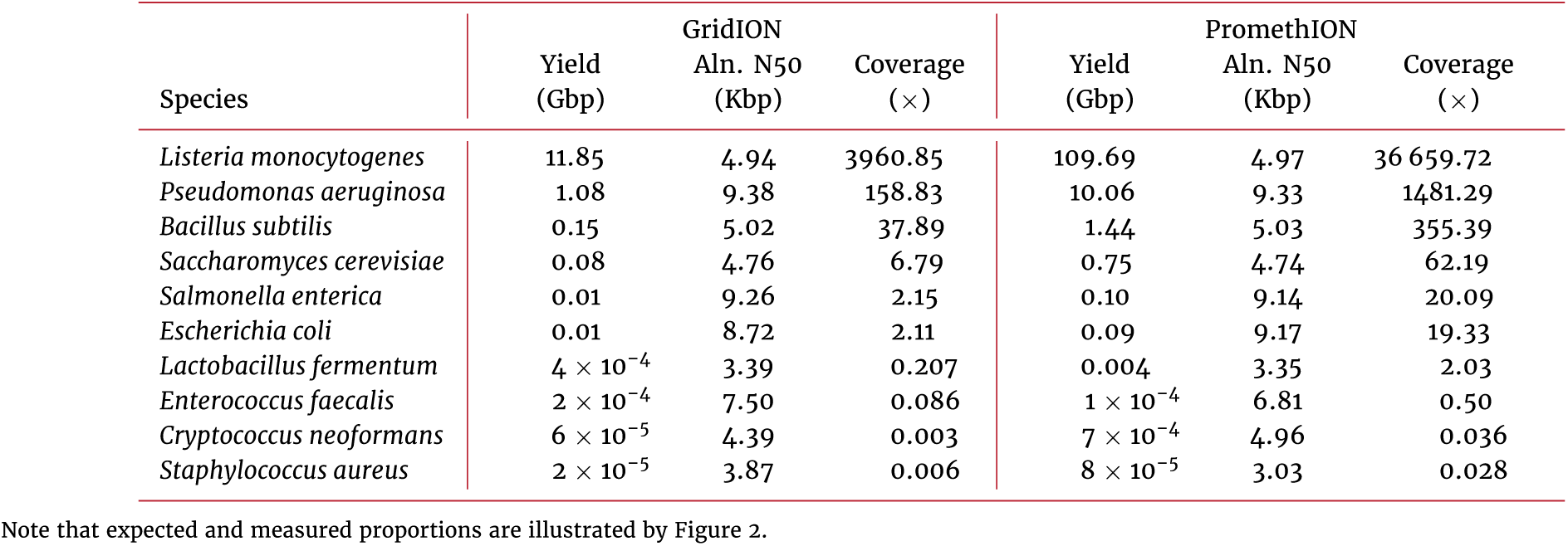
Read alignment statistics for log samples, describing sequencing yield and estimated genome coverage obtained for each organism in the mock community.

**Figure 2.**
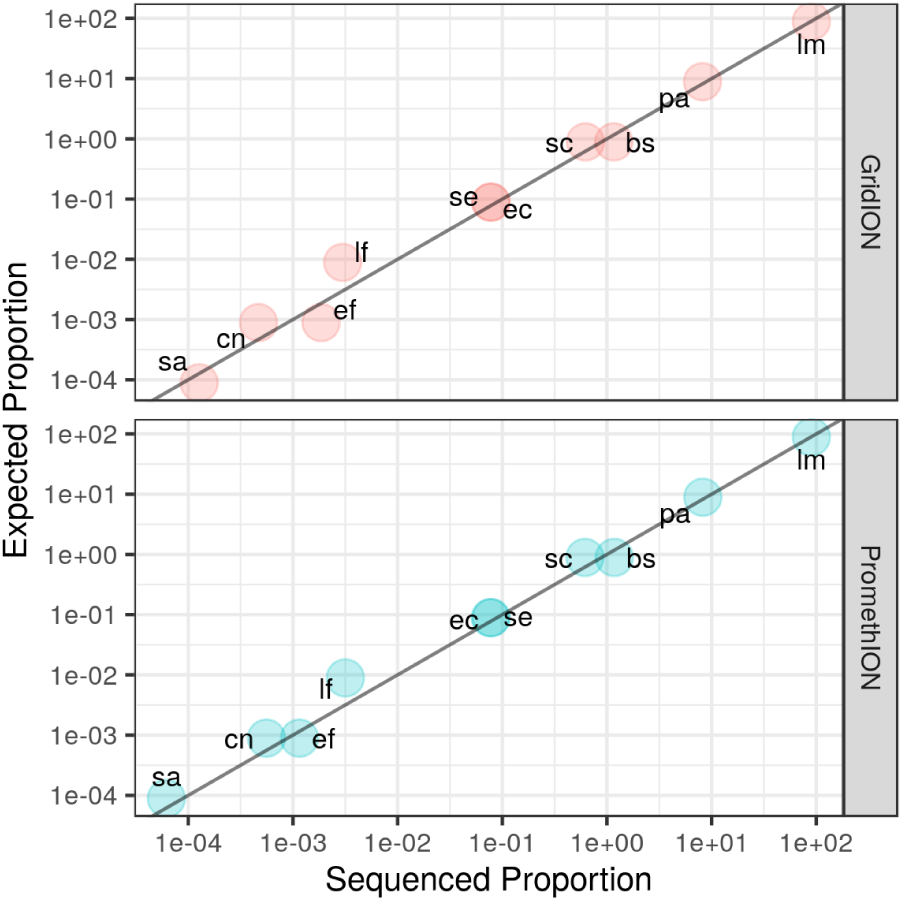
Proportion of sequenced DNA from each of the 10 organisms that was sequenced (x-axis), against the proportion of yield expected given the known community composition (y-axis) of the Zymo CSII (Log) standard.

For the log-distributed community, three taxa have suicient coverage for assembly on GridION, compared with four on PromethION. On PromethION, a further two genomes (*S. enterica* and *E. coli*) have suicient coverage for assembly scaffolding. We are able to detect *S. aureus*, the lowest abundance organism on both platforms, with 32 reads from PromethION (from 450 cell input) and 7 reads from GridION (from 50 cell input).

### Nanopore metagenome assemblies

We evaluated nanopore metagenomic assemblies and assessed genome completeness and contiguity for each run with different assembly parameters.

For the even community, genomes of the expected size were present for each of the bacterial species contained in small numbers of large contigs (Figure 3). However, the two yeasts are poorly represented in the assembly, consistent with their low read depth.

*L. monocytogenes* is poorly assembled in the log dataset despite being the most abundant organism, indicating very high sequence coverage may be detrimental to the performance of `wtdbg2`. We also observe that assembling the entire PromethION dataset resulted in less complete and more fragmented assemblies. This led us to sample randomly the PromethION data to 25% of the total dataset which improved the assembly results.

After subsampling, assemblies of the even community from the GridION and PromethION are similar. However, the assemblies from PromethION data had better representation of the yeasts (particularly *C. neoformans*), due to the higher coverage depths of these species.

Average consensus accuracy was calculated for the entire set of assemblies (n=51). For the even community, accuracy is 99.1% (± 0.17%) and 99.0% (± 0.24%) for the GridION and PromethION respectively. For the log community, average accuracy is 99.0% (± 0.27%) and 99.1% (± 0.20%) for the GridION and PromethION respectively.

## Discussion

There are several noteworthy aspects of this dataset: We generated nearly 300 Gbp of sequence data from the Oxford Nanopore PromethION and 30 gigabases from the Oxford Nanopore GridION, on a well-characterised mock community sample and we have made basecalls and electrical signal data for each of the four runs presented here available: a combined dataset size of over four terabytes. The availability of the raw signal permits future basecalling of the data (an area under rapid development), as well as signal-level polishing and the detection of methylated bases [20].

Individual sequencing libraries were split between the GridION and PromethION, permitting direct comparisons of the instruments to be made. We observed high concordance between the datasets from each platform. We note the sequencing speed of the PromethION is faster than the GridION, which we attribute to different running temperatures on these instrument (39°C versus 34°C, respectively).

Confident detection of *S. aureus* was demonstrated for the GridION run to <50-cells using the log community. The PromethION generated around five times the number of *S. aureus* reads as the GridION, but appears less sensitive as we loaded eight times more library. However, it may be possible to reduce the loading input to PromethION flowcells, but we have not attempted this.

Early results of metagenomics assembly show promise for reconstruction of whole microbial genomes from mixed samples without a binning step. We focused on the recently released `wtdbg2` software as we found that the established `minimap2` and `miniasm` method resulted in excessively large intermediate files (tens of terabases per analysis) which were impractical to store and analyse.

For the even community, using `wtdgb2` with varying parameter choices, we were able to assemble seven of the bacteria into single contigs. However, no single parameter set was found to be optimum for both total genome size and contig length.

We found that `wtdbg2` expects a maximum of 200× sample coverage, and discards sequence k-mers and *de Bruijn* graph nodes with more than 200× support. These limits must be lifted by specifying higher `-K` and `–node-max` for this dataset. Increasing `-e` improved contiguity for the even sample, however this resulted in the loss of yeasts from the assembly. Increasing the read length threshold (`-L`) improved assembly contiguity for all sample and platform combinations, at the cost of genome size. Increasing the homopolymer-compressed k-mer size (`-p`) from the default of 21 to 23 also appears to improve assembly contiguity. It should be noted that `wtdbg2` is still under active development, making it difficult to make concrete recommendations for parameters.

The availability of this dataset should help with further improvements to long-read assembly techniques.

This study has several shortcomings:

We have not yet explored polishing techniques to improve consensus accuracy of assemblies using nanopore or Illumina data [21]. A new “flip-flop” basecaller, `Flappie` https://github.com/nanoporetech/flappie), was recently made available although we have not used it on this dataset.

Although reference genomes for the ten microbial strains contained in the standards are available from Zymo (https://s3.amazonaws.com/zymo-files/BioPool/ZymoBIOMICS.STD.refseq.v2.zip), they are constructed from a combination of nanopore and Illumina data (not presented here). To avoid a circular comparison, we chose to perform analysis only against Illumina draft genomes.

Other mock microbial samples are available which we did not test here. A notable alternative mock community sample is from the Human Microbiome Project (HMP) and consists of 20 microbial samples (available from BEI Resources). This mock community have been sequenced as part of other studies, although the datasets are much smaller than the ones presented here [9, 22]. Bertrand et al. presented a synthetic mock community of their own construction to demonstrate hybrid nanopore-Illumina metagenome assemblies [12].

### Re-use potential

The provision of Illumina reads for each isolate permits a ground-truth to be obtained for the individual species contained in the mock community. This will be useful for training new nanopore basecalling and polishing models, long-read aligners, variant callers, and validating taxonomic assignment and assembly software and pipelines.

## Availability of source code and requirements

Python and R scripts used to generate the summary information and analyses are open source and freely available via our repository (https://github.com/LomanLab/mockcommunity), under the MIT license.

## Availability of supporting data and materials

This manuscript, and its supporting data are available under a Creative Commons Attribution 4.0 International license.

Unprocessed FASTQ from the Illumina sequencing of the ten isolates are available at the European Nucleotide Archive, via the identifiers listed in Table 1, identifiers for the even and log community Illumina sequencing can be found in Table 5.

Both the raw signal, and basecalled FASTQ for the nanopore sequencing experiments are available at the European Nucleotide Archive, via the identifiers listed in Table 2.

The `SPAdes`-assembled Illumina draft reference, and the collection of nanopore assemblies for each `wtdbg2` condition are linked to from our GitHub repository (https://github.com/LomanLab/mockcommunity), along with the `kraken2` database used for taxonomic classification of the assembled contigs.

Further updates (such as updated references, or new assemblies) will be made available through our project website https://lomanlab.github.io/mockcommunity/.

## Declarations

### Consent for publication

Not applicable

### Author’s Contributions

Conceptualization: NL, Methodology: NL JQ SN ST, Software: SN NL, Validation: SN NL, Formal analysis: SN NL, Investigation: NL JQ SN, Resources: NL ST, Data Curation: SN NL ST, Writing – original draft preparation: SN, Writing – review and editing: SN NL JQ ST, Visualization: SN NL, Supervision: NL, Project administration: NL, Funding acquisition: NL ST

## Acknowledgements

We are grateful to Radoslaw Poplawski (University of Birmingham) for assistance with CLIMB virtual machines and file systems to support this research. We thank Divya Mirrington (Oxford Nanopore Technologies) for advice on PromethION library preparation and sequencing. We thank Hannah McDonnell at Cambridge Biosciences for providing the ZymoBIOMICS Microbial Community Standards. We thank Jared Simpson (Ontario Institute for Cancer Research), Matt Loose (University of Nottingham) and John Tyson (University of British Columbia) for useful discussions and advice.

